# Phosphorus acquisition efficiency in arbuscular mycorrhizal maize is correlated with the abundance of root-external hyphae and the accumulation of transcripts encoding PHT1 phosphate transporters

**DOI:** 10.1101/042028

**Authors:** Ruairidh J. H. Sawers, Simon F. Svane, Clement Quan, Mette Grønlund, Barbara Wozniak, Eliécer González-Muñoz, Ricardo A. Chávez Montes, Ivan Baxter, Jerome Goudet, Iver Jakobsen, Uta Paszkowski

## Abstract

- Plant interactions with arbuscular mycorrhizal fungi have long attracted interest for their potential to promote more efficient use of mineral resources in agriculture. Their widespread use, however, remains limited by understanding of the processes that determine the outcome of the symbiosis. In this study, variation in growth response to mycorrhizal inoculation was characterized in a panel of diverse maize lines.
- A panel of thirty maize lines was evaluated with and without inoculation with arbuscular mycorrhizal fungi. The line Oh43 was identified to show superior response and, along with five other reference lines, was characterized in greater detail in a split-compartment system, using ^33^P to quantify mycorrhizal phosphorus uptake.
- Changes in relative growth between non-inoculated and inoculated plants indicated variation in host capacity to profit from the symbiosis. Shoot phosphate content, abundance of intra-radical and root-external fungal structures, mycorrhizal phosphorus uptake, and accumulation of transcripts encoding plant PHT1 family phosphate transporters varied among lines.
- Larger growth responses in Oh43 were correlated with extensive development of root-external hyphae, accumulation of specific *Pht1* transcripts and a high level of mycorrhizal phosphorus uptake. The data indicate that host genetic factors influence fungal growth strategy with an impact on plant performance.

## INTRODUCTION

The rising cost of agricultural inputs and an increasing awareness of the potential negative environmental consequences of their use have resulted in ever greater interest in beneficial crop-microbe interactions and their application (Perez-Montano *et al.*, 2014; Vance, 2014). The most prevalent nutrient-delivering symbiosis is the association of plants with fungi of the phylum *Glomeromycota*, resulting in the formation of arbuscular mycorrhizas (Smith & Read, 2008). More than 80% of extant terrestrial plants establish arbuscular mycorrhizal (AM) symbioses, and this fundamental capacity has been retained in major crop species throughout the processes of domestication and improvement (*e.g.* Koide *et al.*, 1988; Hetrick *et al.*, 1992; Kaeppler *et al.*, 2000; Sawers *et al.*, 2008). Concomitantly, these same crops have retained a conserved molecular machinery required for symbiotic establishment and nutrient exchange (Paszkowski *et al.*, 2002; Gutjahr *et al.*, 2008; Yang *et al.*, 2012; Willman *et al*., 2013).

The best characterized benefit of AM symbiosis is enhanced plant phosphorus (P) nutrition. The efficiency with which crop plants convert P resources to yield (P Efficiency; PE) can be partitioned between the efficiency of uptake (P Acquisition Efficiency; PAE), and the efficiency of internal use (P Use Efficiency; PUE) (Rose *et al*., 2011; Veneklaas *et al.*, 2012). Plants take up P in the form of phosphate (Pi), and PAE is typically low in agricultural systems (15-20% of applied P taken up by plants; Syers *et al.*, 2008) as the result of the low mobility of Pi in the soil and the ready formation of a zone of P depletion around the root. By extending beyond the P depletion zone, the root-external hyphae of AM fungi increase the extent of soil foraging and P uptake (Bucher, 2007). Physiological studies have demonstrated that Pi uptake via AM fungi is a distinct functional alternative to direct uptake by the plant (Smith *et al.*, 2003; Bucher, 2007), and, in a field setting, the majority of the Pi taken up by a plant may be acquired via the mycorrhizal route (Schweiger and Jakobsen, 1999; Smith *et al.*, 2003; Yang *et al.*, 2012). Molecular analyses support the distinction between mycorrhizal and direct P uptake, members of the plant PHT1 P transporter family having been identified to play roles specific to the two pathways (Bucher, 2007). Where plants are competent to host AM fungi, there is at least one family member acting predominantly, or exclusively, during AM symbiosis (ZmPT6 in maize and variously named orthologs in other species; here, the short form PT is used to refer to all PHT1 genes and proteins; Rausch *et al.*, 2001; Harrison *et al.*, 2002; Paszkowski *et al.*, 2002; Glassop *et al.*, 2005; Nagy *et al.*, 2005; Maeda *et al.*, 2006; Caesar *et al.*, 2014; Walder *et al.*, 2015). In *Medicago truncatula*, the ZmPT6 ortholog MtPT4 has been localized to the peri-arbuscular membrane (Harrison *et al.*, 2002; Kobae & Hata, 2010; Pumplin *et al.*, 2012), consistent with a function as the principal route of P uptake from fungus to plant.

While AM symbiosis can increase PAE and also provide enhanced tolerance to a range of abiotic and biotic stresses (Smith & Read, 2008), such benefits are not provided without cost. The plant host must supply carbohydrates to support fungal growth, diverting photosynthetically fixed carbon away from primary productivity. Ultimately, the outcome, in terms of the impact on plant growth and fitness (or yield in crop plants), will depend not only on the specific plant-fungus combination (Walder *et al.*, 2012) but on the requirements and limitations imposed by any given environment (Janos, 2007). In high-input agricultural systems, AM symbiosis may provide no benefit to yield, or even be detrimental (Grace *et al.*, 2009), and it has been hypothesized that plant breeding may have promoted a weakening of the mutualism in modern elite cultivars (Hetrick *et al.*, 1992, 1996). Drawing firm conclusions as to impact of AM symbiosis, however, is difficult. Mycorrhizal response (defined here as MR = M – NC, where M is the trait value for colonized plants and NC the trait value for non-colonized plants) confounds plant adaptation to a given set of conditions and capacity to benefit from the symbiosis *per se* (discussed in Sawers *et al*., 2010). To emphasize this distinction, the term dependence has been defined as the nutrient requirement of a particular variety to attain a certain threshold level of performance (Janos *et al.*, 2007). In practice, poorly adapted (*i.e.* highly dependent) varieties will typically show a large performance increase following AM colonization (e.g. Hetrick *et al*., 1992; Kaeppler *et al*., 2000; Paszkowski & Boller, 2002): such varieties, although highly responsive, are clearly of little agronomic interest. The question remains as to whether certain varieties derive greater benefit from AM symbioses *per se* than others and to what extent plant breeding can optimize these interactions for agricultural systems (Sawers *et al.*, 2008; Fester & Sawers, 2011).

The objective of this study was to identify maize line(s) highly responsive to mycorrhiza inoculation, conducting detailed physiological analysis to demonstrate greater benefit in selected line(s) when colonized.

## MATERIALS AND METHODS

### Evaluation of response to *Funneliformis mosseae* in maize diversity panel

A panel of 30 diverse maize lines, comprising the 26 diverse inbred founders of the maize NAM population (McMullen *et al.*, 2009), Pa36 (a line tolerant of low P availability; Kaeppler *et al.*, 2000), and the broadly used reference lines B73 and W22, and W64A (a line used previously for study of AM symbiosis; Paszkowski *et al.*, 2006), was evaluated with (M) or without (NC) inoculation with *F. mosseae* (isolate number 12, European Bank of Glomales, http://www.kent.ac.uk/bio/beg/). Plants were grown in 1 litre pots in a mixture of 1:10 loam: quartz sand, as previously described (Sawers *et al.*, 2010). The greenhouse was maintained at 28°C day/night temperature with a 12-h light period, including supplementary light when required. Plants were fertilized three times per week with 100 ml of modified Hoagland solution (Hoagland and Broyer, 1936) containing 10% (100μM) of the standard concentration of KH_2_PO_4_, the potassium concentration being maintained by addition of KCl. A total of 1200 plants (30 genotypes x 2 treatments x 20 replicates) were grown in a complete block design, over five separate plantings, at the University of Lausanne, Switzerland. Plants were harvest 8 weeks and shoot dry weight measured (SDW). For two plantings (corresponding to six complete blocks) roots were collected also, and stained to confirm efficacy of the fungal inoculum as previously described (Gutjahr *et al.*, 2008). Prior to analysis of SDW, data was adjusted to account for differences among plantings. The main effect of inoculation (mycorrhiza response (MR) = M – NC; Sawers *et al*., 2010) was estimated as the difference in means of M of NC groups and shown to be significant using a *t-*test (p < 0.001). MR was estimated similarly for each genotype and a 95% *t*-interval calculated for the difference M – NC. ANOVA (R statistics; R Core Team, 2016) was used to compare SDW among the sixty line/treatment combinations (p <0.001) and means groups were calculated using least significant difference (R statistics agricolae::LSD.test; de Mendiburu, 2016).

### Determination of elemental concentration by ICP-MS analysis

Weighed tissue samples were digested in 2.5mL concentrated nitric acid (AR Select Grade, VWR) with internal standard added (20ppb In, BDH Aristar Plus). Sample digestion and dilution was carried out as described previously (Ziegler *et al.*, 2013). Elemental concentrations of B, Na, Mg, Al, P, S, K, Ca, Mn, Fe, Co, Ni, Cu, Zn, As, Se, Rb, Mo, and Cd were measured using an Elan 6000 DRC-e mass spectrometer (Perkin-Elmer SCIEX) connected to a PFA microflow nebulizer (Elemental Scientific) and Apex HF desolvator (Elemental Scientific). A control solution was run every tenth sample to correct for machine drift both during a single run and between runs. Statistical analysis was conducted with R statistics (see above).

### Characterization of AM phosphorus uptake in six selected lines

Six maize lines, selected on the basis of pre-screening of a panel of 30 lines, were evaluated at the Technical University of Denmark. Plants were grown in 2.4 L PVC tubes in accordance with Smith *et al.* (2003). The growth medium (hereafter referred to as soil) was a 1:1 (w:w) mixture of quartz sand and γ-irradiated soil (15 kGy) that had basal nutrients (Pearson & Jakobsen, 1993) and KH_2_PO_4_ at nil, 15 or 90 mg P kg^−1^ added in solution and carefully mixed into the soil. The P additions resulted in a bicarbonate-extractable P content of 7.9, 15.5 or 53.3 mg P kg^−1^ (Olsen *et al.*, 1954). The root/hyphae compartment (RHC) contained 2750 g soil and the hyphae compartment (HC) was a small plastic vial placed in the middle of the RHC. The HC contained 55 g of ^33^P-labeled soil (5 kBq g^−1^) and was lined with a 25 μm nylon mesh at both ends to prevent root in-growth. Seven weeks later, bicarbonate extracts of HC had a specific activity (SA = ^33^P/^31^P) of 144.7, 79.9 or 29.4 kBq mg^−1^ P, corresponding to no addition or addition of 15 or 90 mg P kg^−1^. Each maize line was grown in 8 replicate pots in half of which 140 g dry soil-root inoculum of *Rhizophagus irregularis* BEG87 was thoroughly mixed into the growth medium. Filtered BEG87 inoculum leachings were added to all pots as an attempt to establish the same soil microbial community (Pearson & Jakobsen, 1993). Two pre-germinated seeds were planted in each pot and thinned to one at the two leaf stage. Plants were maintained under controlled conditions (12 hour day length at 500 μmol m^−2^ sec^−1^, 28/20°C day/night and 60 % relative humidity) and watered daily by weight to 70% of the water holding capacity. Plants received supplemental N (NH_4_NO_3_), Mg and S (MgSO_4_ ^2-^) periodically to additionally provide 375 mg N, 15 mg Mg and 20 mg S per pot. Shoots were harvested at growth stage 51 (BBCH scale; tassel emergence at the top of the stem), oven dried to constant weight at 70°C and dry weights were recorded. Root systems were carefully washed clean using a pressurized water jet and a fine mesh to collect fine root pieces. Roots were blotted dry and total fresh weight (FW) was recorded. Subsamples were taken for root length/colonization measurement (1.5g FW, stored in 50% EtOH) and RNA extraction (1g, flash-frozen in liquid nitrogen). Dried and ground shoot and root samples were oxidized in a 4:1 mixture (v:v) of 65% nitric:70% perchloric acids, and total P was determined by the molybdate blue method using AutoAnalyzer 3 (Bran+Luebbe, Norderstedt, Germany). The ^33^P in shoot tissue was determined in the same digests in a Packard TR 1900 liquid scintillation counter (PerkinElmer, Waltham, MA, USA). Specific activities of ^33^P in shoots and the specific activities in bicarbonate extracts of HC soil were used to estimate the amount of P taken up from the HC in accordance with Smith et al. (2004). The abundance of total fungal structures (hyphae, arbuscules or vesicles) or arbuscules specifically was evaluated microscopically as percentage of root length using the grid-line intersect method (Newman, 1966) after clearing and staining (Kormanik & McGraw, 1982). The length of hyphae in HC soil was measured by an aqueous extraction and membrane filter technique (Jakobsen *et al.*, 1992) with the modification that the stained filters were mounted on slides using immersion oil instead of lactoglycerol-trypan blue solution in order to facilitate discrimination of AM and non-AM fungal hyphae. Statistical analysis was conducted with R statistics (see above).

### Bioinformatic identification of maize *Pht* genes

To identify a complete set of putative PHT1 encoding genes in maize, the *Saccharomyces cerevisiae* PHO84 protein (Uniprot id P25297) was used as a BlastP query (Altschul *et al.*, 1990) to search the primary transcript predicted protein sequences from version 6a of the annotated B73 maize genome (Schnable *et al.*, 2009), obtained from Phytozome 10 (Goodstein *et al.* 2012). Using a cut-off E-value of 1e^−54^, 13 gene-models were retrieved and aligned using MUSCLE (Edgar, 2004). All 13 sequences contained the conserved GGDYPLSATIxSE motif in helix 4 reported previously to be present in PHT proteins (Karandashov & Bucher, 2005). The resulting block-alignment file was converted to Stockholm 1.0 format, and used as input to hmmbuild (HMMER suite version 3.1b2) to search (hmmsearch) the maize primary transcript predicted protein sequences for additional PHT1 proteins. 35 new protein sequences were identified based on an inclusion threshold of E-value <0.01. None of these additional sequences, however, contained the conserved GGDYPLSATIxSE motif and consequently there were not considered to be authentic PHT1 proteins. The final list of 13 maize PHT1 genes was consistent with the report of Liu *et al.*, 2016.

### Analysis of *ZmPt* transcript accumulation

Gene specific primers for real time PCR analysis of *ZmPt* transcript accumulation were designed and successfully optimized for twelve of thirteen annotated genes (Table S2). All primer sets were used in a preliminary characterization of B73 plants, and five sets (*Pt3*, *Pt1*, *Pt6*, *Pt4*, *Pt5*,) selected on the basis of accumulation pattern for analysis of accumulation in the roots of six maize lines grown in the P uptake experiment. Samples were prepared using the LightCycler 480 SYBR green I master mix kit (Roche; Mannheim, Germany) before analysis on a Roche 480 LightCycler. Each biological sample was analysed as three technical replicates. Three water controls were used for each gene tested. qRT-PCR expression and melting curves were calculated using the LightCycler 480 software (Roche, Version 1.5.9, Mannheim, Germany). Samples were normalized to the geometric mean of expression levels of 3 constitutive genes (*GAPDH, Cyclophilin2, ß-actin*) as described earlier (Gutjahr *et al.*, 2008). In addition to phosphate transporters were analysed together with an AM specific marker gene *ZmAm3*, ortholog of *OsAM3* (Gutjahr *et al.*, 2008) and a *Rhizophagus irregularis* elongation factor gene (Sokolski *et al.*, 2010). Statistical analysis was conducted with R statistics (see above).

### Principal component analysis of combined growth, physiology and molecular data sets

Pairwise correlations were calculated for a matrix of growth, physiological and molecular data obtained from six selected lines as described above (shoot dry weight, shoot P content, root dry weight, root P content, total colonization, arbuscule abundance, length of root-external hyphae, P uptake from the hyphal compartment, accumulation of *Pt3*, *Pt1*, *Pt6*, *Pt4*, *Pt5*, *Am3* and *RiEF* transcripts) using R statistics Hmisc::rcorr (Harrel FE, 2016)and the results visualized using R statistics corrplot::corrplot (Wei & Simko, 2016). Principal component analysis (PCA) was performed with R statistics ade4::dudi.pca (Dray & Dufour, 2007) using centered and scaled data, and the results visualized with ade4::scatter.

## RESULTS

### The line Oh43 exhibits typical dependence but high mycorrhizal responsiveness

To identify lines showing a typical level of dependence (defined here as growth in the absence of inoculation) but high mycorrhiza response (MR), a panel of thirty diverse maize lines (the parents of the maize Nested Association Mapping population and a number of additional lines; see Materials and Methods) was evaluated with (M) or without (NC) inoculation with the fungus *Funneliformis mosseae*. The composition of the diversity panel was determined previously to maximize sampling of genetic diversity from global maize breeding germplasm (McMullen *et al*., 2009). Use of this panel was designed to provide a context to evaluate the relative performance of any given line, whether inoculated or not.

A total of 1200 plants (30 genotypes x 2 treatments x 20 replicates) were grown in a complete block design, using 1L pots, in P limiting conditions, for a period of eight weeks after emergence (V8 stage). Roots were harvested from 30% of the total experiment and fungal structures by microscopic inspection to confirm the efficacy of inoculation (*Supporting Information*). NC plants were confirmed to be free of fungal structures, while in M plants, the level of colonization was generally high, with a mean of 57% ±0.7% (95% interval for proportion) of root positions examined containing at least one type of fungal structure (hyphae, arbuscules or vesicles), although a broad range of colonization was observed (5% - 98%). For all experimental blocks, the aerial portion of the plants was dried and shoot dry weight (SDW; g) determined (Table 1). Overall, the evaluated lines showed greater growth when colonized by *F. mosseae* (Fig. S1), with a significant (p < 0.001) increase in mean SDW from 1.05g in NC plants to 2.16 g in M plants, equating to panel-wide MR of 1.1 ±0.08 g (MR = M – NC; 95% interval for difference in means). At the level of the individual lines, all showed a significant increase in SDW following inoculation (Fig. 1a; Table 1). The most responsive line Oh43 showed a significantly greater MR than the least responsive line Mo18W (MR_Oh43_ = 1.85 ±0.54 g; MR_Mo18W_ = 0.72 ±0.28 g; 95% t-intervals for MR do not overlap). The contrast in outcome between Oh43 and Mo18W was reflected by rank-changing shifts in SDW relative to other lines in the panel between NC and M conditions: Oh43 and Mo18W were similar to each other, and typical of the panel as a whole, when not inoculated, but differed to each other and were outlying when inoculated (Fig. 1a, b; Table 1; LSD, α = 0.05). The line Oh43, by exhibiting typical dependence but high MR, fulfilled the criteria established to identify superior capacity to benefit from AM symbiosis.

**Table 1.** Shoot dry weight of 30 maize lines grown with or without inoculation with *Funneliformis mosseae.* Shoot dry weight of 8 week-old non-colonized (NC; *n* = sample size) and mycorrhizal (M; inoculated with *F. mosseae*; *n* = sample size) plants. Means groups based on least significant difference (LSD) were calculated separately for NC and M plants at α = 0.05. Mycorrhiza response (MR) was calculated as M-NC, numbers in parentheses give a 95% confidence interval (CI) for the difference in means.

| Line | NC (g) | $n$ | NC LSD | M (g) | $n$ | M LSD | MR (95% CI) |
| --- | --- | --- | --- | --- | --- | --- | --- |
| Oh43 | 0.96 | 20 | ghijklm | 2.81 | 20 | ab | 1.85 (1.31 ,2.38) |
| CML322 | 1.13 | 18 | efghi | 2.59 | 17 | abcde | 1.47 (0.8 ,2.14) |
| HP301 | 0.44 | 18 | n | 1.82 | 18 | ij | 1.37 (0.91 ,1.84) |
| A188 | 0.98 | 13 | fghijklm | 2.3 | 14 | bcdefghi | 1.32 (1.04 ,1.61) |
| W64 | 1.02 | 20 | fghijkl | 2.34 | 19 | bcdefgh | 1.32 (0.91 ,1.73) |
| B97 | 0.74 | 20 | m | 2.04 | 20 | fghij | 1.31 (0.89 ,1.73) |
| NC350 | 0.86 | 18 | jklm | 2.17 | 18 | defghi | 1.31 (0.8 ,1.81) |
| M162W | 1 | 17 | fghijkl | 2.23 | 14 | cdefghi | 1.23 (0.86 ,1.61) |
| P39GB | 0.87 | 18 | ijklm | 2.11 | 19 | efghij | 1.23 (0.79 ,1.67) |
| Pa36 | 1.67 | 15 | a | 2.91 | 15 | a | 1.23 (0.71 ,1.76) |
| Ms71 | 1.4 | 19 | bcd | 2.62 | 20 | abcd | 1.22 (0.71 ,1.73) |
| CML333 | 1.19 | 20 | defg | 2.4 | 20 | abcdefg | 1.2 (0.8 ,1.61) |
| Mo17 | 1.53 | 17 | ab | 2.7 | 20 | abc | 1.17 (0.7 ,1.64) |
| CML103 | 1.37 | 19 | bcde | 2.52 | 20 | abcdef | 1.15 (0.87 ,1.43) |
| CML52 | 0.81 | 19 | klm | 1.9 | 19 | hij | 1.1 (0.7 ,1.49) |
| Ki11 | 1.02 | 19 | fghijkl | 2.1 | 20 | efghij | 1.08 (0.75 ,1.42) |
| IL14H | 0.96 | 14 | fghijklm | 2.03 | 15 | fghij | 1.07 (0.8 ,1.34) |
| B73 | 0.78 | 16 | lm | 1.82 | 17 | ij | 1.05 (0.78 ,1.31) |
| CML277 | 0.93 | 20 | hijklm | 1.96 | 20 | ghij | 1.03 (0.6 ,1.45) |
| CML228 | 1.38 | 19 | bcd | 2.35 | 20 | bcdefgh | 0.97 (0.66 ,1.27) |
| M37W | 0.94 | 20 | hijklm | 1.91 | 19 | hij | 0.97 (0.38 ,1.57) |
| CML247 | 1.08 | 18 | fghij | 1.98 | 20 | ghij | 0.9 (0.62 ,1.18) |
| Tx303 | 0.7 | 18 | mn | 1.61 | 17 | j | 0.9 (0.46 ,1.34) |
| Ki3 | 0.94 | 20 | hijklm | 1.79 | 20 | ij | 0.85 (0.53 ,1.16) |
| Ky21 | 1.23 | 18 | cdef | 2.08 | 18 | fghij | 0.84 (0.49 ,1.19) |
| NC358 | 1.22 | 17 | cdef | 2.03 | 18 | fghij | 0.8 (0.42 ,1.19) |
| W22 | 1.15 | 17 | defgh | 1.93 | 16 | ghij | 0.78 (0.43 ,1.13) |
| Oh7b | 1.05 | 20 | fghijk | 1.81 | 20 | ij | 0.76 (0.39 ,1.13) |
| Tzi8 | 1.48 | 15 | abc | 2.24 | 19 | cdefghi | 0.76 (0.38 ,1.15) |
| Mo18W | 0.94 | 18 | hijklm | 1.66 | 20 | j | 0.72 (0.44, 1) |

**Fig. 1.**
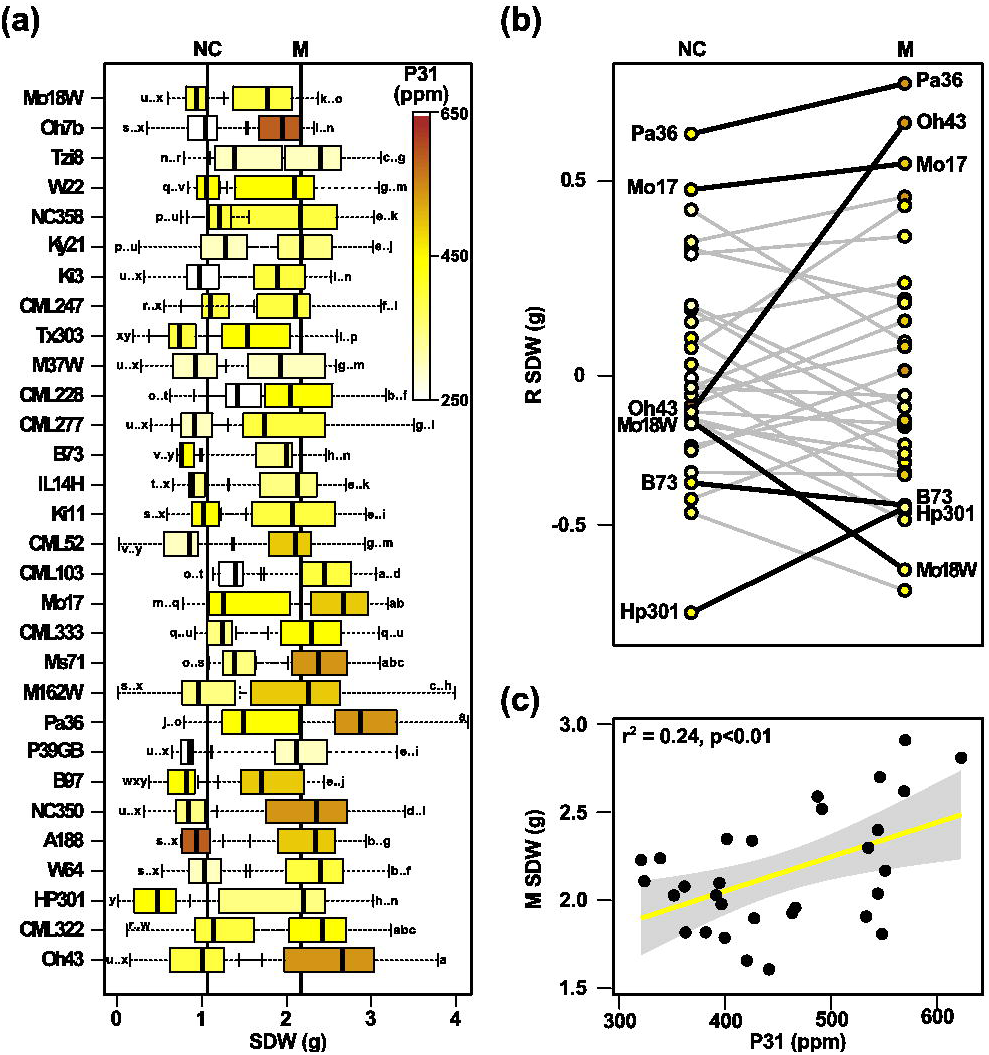
Maize lines varied in response to inoculation with *Funneliformis mosseae*. (a) Shoot dry weight (SDW, g; normalized with respect to differences among experimental plantings) of 30 diverse maize lines grown for 8 weeks with (M; right box) or without (NC left box) inoculation with the fungus *F. mosseae*. Boxes show 1st quartile, median and 3rd quartile. Whiskers extend to the most extreme points within 1.5x box length; outlying values beyond this range are not shown. The overall mean values of NC (1.05g, n=540) and M (2.16g, n=552) groups are shown by vertical lines. Lines are ordered by increasing mycorrhizal response (M-NC) from top to bottom. Letters adjacent to boxes indicate means groups, calculated for NC and M treatments collectively on the basis of least significant difference at α = 0.05. Box shading indicates mean phosphorus content (P^31^, ppm) in the shoot as determined by ionomic analysis, colour-key shown at the top of the panel. (b) Reaction norms (plot of phenotype against environment) for 30 diverse maize lines, contrasting shoot dry-weight (R SDW, g; residual SDW with respect to overall NC or M mean) of non-inoculated plants (NC) and plants inoculated with *F. mosseae* (M). Segments corresponding to six lines selected for further study are labeled and shown in bold. Point shading indicates shoot P content (ppm) as (a). (c) Shoot dry weight (SDW, g) as a function of shoot P content (P^31^, ppm) in 30 maize lines inoculated with *F. mosseae* (points correspond to mean values). Linear fit (yellow line) and associated 95% confidence interval (shaded area) shown.

Evaluation was conducted under P limiting conditions, and it was expected that variation in growth would be driven largely by differences in P efficiency. Leaf P concentration (ppm) was quantified using inductively coupled plasma-mass spectroscopy (Table S1) and correlated positively with SDW in M plants (Fig. 1c; p<0.01, r^2^ = 0.24). Leaf P concentration was similar in Mo18W and Oh43 when non-colonized but different when plants were inoculated (Fig. 1a, b; Table S1; LSD, α = 0.05), mirroring SDW, and suggesting the difference in MR between the two lines to be the result of greater PAE in Oh43 when colonized.

In summary, evaluation of shoot dry weight and P concentration in the diversity panel identified the line Oh43 to be highly responsive to inoculation with *F. mosseae* on the basis of functional differences in the symbiosis.

### High mycorrhizal responsiveness in Oh43 is correlated with abundant root-external hyphae

To characterize further the mechanistic basis of MR in Oh43, the line was evaluated in a previously described split-compartment system, separating root/hyphae (RHC) and hyphae compartments (HC) for direct quantification of AM mediated P uptake through ^33^P labeling (Smith *et al.*, 2003). Although it was not feasible to evaluate the complete panel in this more detailed study, five additional lines were included to provide a context for interpretation of the data, namely the low MR line Mo18W, the widely used reference lines B73 and Mo17, and HP301 and Pa36, the lines that showed the highest and lowest NC SDW, respectively, in the panel evaluation (Fig. 1; Table 1). Eight replicates of the six lines were evaluated at three P levels (low = 7.9 mg P kg^−1^; medium = 15.5 mg P kg^−1^; high = 53.2 mg P kg^−1^), with (M) or without (NC) inoculation with *Rhizophagus irregularis*, a fungal species that has been previously used with success in this system. Given that plants may respond differently to different fungal species and strains (*e.g.* Angelard *et al.*, 2010), the following discussion and further conclusions are confined to this *R. irregularis* experiment.

Plants were harvested at tassel emergence, and samples taken for measurement of SDW, P content, and abundance of fungal structures. In addition, soil was collected from the HC for quantification or root-external hyphae. In all lines, SDW increased with greater P addition, irrespective of inoculation status (Fig. 2; Table2). In contrast, MR decreased from positive to negative with increasing P availability (mean of six lines: low P, mean MR = 9.56g; medium P, mean MR = 3.95g, high P, mean MR = −1.57g), the ranking of the lines with respect to MR changing also. At low P, Oh43 was more responsive than the other lines, consistent with the results of the *Funneliformis mosseae* experiment, and remained highly responsive at medium P, although exhibiting negative MR at high P (Fig. 2). As observed previously in the *F. mosseae* experiment, SDW was correlated with shoot P content (Fig. 2). Under low P, shoot P content in Oh43 was typical in non-inoculated plants, but high when plants were colonized. Quantification of P uptake from the HC indicated that Oh43 was indeed obtaining more P via the mycorrhizal pathway under low P than the other lines (Table 2). When the roots were examined, however, Oh43 was not more heavily colonized; indeed, the proportion of root length containing arbuscules was marginally lower (Fig. 3a). was observed to be significantly greater than in the other lines (Fig. 3b; Table 2). Among the six lines, and across all three levels of P availability, a clear relationship was observed between P uptake from the HC and the length of root-external hyphae (Fig. 3b). At high P, the growth response was apparently saturated in the lines B73 and Pa36, which attained their maximum growth irrespective of AM inoculation, although P content was lower in the shoots of M plants than NC plants (*i.e.* PUE was greater in M plants). Interestingly, although inoculated B73 and Pa36 plants showed equivalent growth, P uptake and abundance of intra-radical fungal structures at high P, the length of root-external hyphae was greater in B73. Indeed, B73 supported a generally high level of root-external hyphae: second only to Oh43 at low P, and greater than all other lines at high P. Collectively, these data indicate that Oh43 is highly responsive to inoculation with *R. irregularis* at low P availability, with concomitant extensive development of root-external hyphae and high PAE in inoculated plants.

**Fig. 2.**
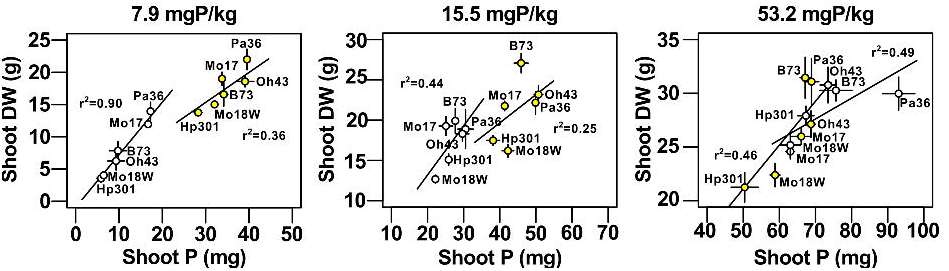
Plant growth was correlated with phosphorus content in six maize lines inoculated with *Rhizophagus irregularis*. Shoot dry weight (Shoot DW; g) as a function of total phosphorus (P) content (Shoot P; mg) in six maize genotypes grown with (yellow circles) or without (open circles) inoculation with *R. irregularis*, under three levels of P availability (7.9, 15.5, 53.2 mgP/kg). Points indicate the mean; whiskers extend +/- 1 standard error; trend lines based on a linear fit to individual observations for each treatment.

**Table 2.** Characterization of six maize lines grown with or without inoculation with *Rhizophagus irregularis* and at three P levels. Shoot dry weight (SDW) and shoot P content (SP) of non-colonized (NC) and mycorrhizal (M) plants; Phosphorus obtained from the hyphae compartment (PHC) in M plants; % of root length containing fungal structures in M plants (Col); % of root length containing arbuscules in M plants (Arb); Hyphal length density in the hyphae compartment in M plants (HLD). Numbers are means and standard errors (n=4). Means groups based on least significant difference were calculated separately for each P level. For SDW and SP, means groups were calculated for NC and M plants together. For Col and Arb, means groups were calculated using square root transformed data.

| Line | Shoot dry wt (g) |  | Shoot P content (mg) |  | P from HC (mg) | Col % | Arb % | HLD (m/g) |
| --- | --- | --- | --- | --- | --- | --- | --- | --- |
|  | NC | M | NC | M |  |  |  |  |
| 7.9 mg P kg<sup>-1</sup> |  |  |  |  |  |  |  |  |
| Oh43 | 6.3 (1.7) ef | 18.6 (1.0) b | 9.3 (2.3) ef | 39.1 (2.3) a | 0.54 (0.06) a | 87.8 c | 70.4 b | 21.4 (2.5) a |
| Mo18W | 4.0 (0.2) f | 15.0 (0.5) cd | 6.5 (0.4) fg | 32.1 (0.2) b | 0.34 (0.05) b | 94.8 a | 79.1 a | 13.4 (0.8) bc |
| Pa36 | 14.0 (1.6) cd | 22.0 (1.9) a | 17.3 (0.6) d | 39.5 (0.8) a | 0.41 (0.06) b | 93.9 ab | 83.4 a | 16.3 (2.9) b |
| Hp301 | 3.6 (0.7) f | 13.8 (0.6) cd | 5.9 (1.2) g | 28.3 (1.1) c | 0.20 (0.02) c | 92.6 abc | 77.9 ab | 10.5 (0.7) c |
| B73 | 7.8 (1.6) e | 16.6 (2.1) bc | 9.8 (2.3) e | 34.2 (1.1) b | 0.41 (0.04) b | 92.5 abc | 84.6 a | 17.0 (2.2) ab |
| Mo17 | 12.0 (0.5) d | 19.0 (1.3) ab | 16.7 (0.4) d | 33.8 (0.9) b | 0.41 (0.02) b | 88.0 bc | 78.7 ab | 13.0 (0.8) bc |
| 15.5 mg P kg<sup>-1</sup> |  |  |  |  |  |  |  |  |
| Oh43 | 18.3 (1.0) def | 23.2 (1.3) b | 29.7 (2.1) de | 50.7 (1.4) a | 0.83 (0.14) ab | 89.5 bc | 65.5 c | 19.4 (3.1) a |
| Mo18W | 12.7 (0.6) g | 16.2 (0.9) ef | 22.2 (1.0) f | 42.3 (1.9) bc | 0.68 (0.07) bc | 96.8 a | 82.1 a | 14.7 (1.3) bc |
| Pa36 | 18.9 (2.7) cde | 22.1 (1.6) bc | 30.5 (2.4) d | 49.9 (4.4) a | 0.72 (0.1) abc | 96.6 a | 78.5 ab | 14.2 (0.7) c |
| Hp301 | 15 (0.8) fg | 17.5 (0.7) def | 25.9 (0.6) def | 38.1 (2.1) c | 0.54 (0.1) c | 91.7 b | 76.8 ab | 13.2 (1.5) c |
| B73 | 19.9 (1.8) bcd | 27.1 (1.4) a | 27.7 (1.0) de | 45.9 (2.2) ab | 0.92 (0.03) a | 93.0 ab | 72.9 b | 18.9 (2.2) ab |
| Mo17 | 19.3 (1.4) cde | 21.8 (0.7) bc | 25.1 (2.2) ef | 41.4 (0.6) bc | 0.76 (0.1) ab | 87.9 c | 71.8 c | 15.0 (0.8) abc |
| 53.2 mg P kg<sup>-1</sup> |  |  |  |  |  |  |  |  |
| Oh43 | 30.8 (1.9) ab | 27.1 (1.4) bc | 73.5 (2.5) bc | 68.8 (1.3) bcd | 0.69 (0.1) b | 74.7 b | 45.0 a | 13.9 (1.3) ab |
| Mo18W | 25.2 (1.3) cd | 22.4 (1.3) de | 63.0 (3.3) de | 58.8 (1.6) e | 0.69 (0.03) b | 84.0 a | 55.7 a | 10.2 (0.8) bc |
| Pa36 | 30.0 (1.8) ab | 31.1 (2.5) a | 93.0 (5.2) a | 68.9 (2.4) bcd | 0.81 (0.2) b | 87.7 a | 54.1 a | 10.5 (1.4) abc |
| Hp301 | 27.9 (1.0) abc | 21.3 (1.6) e | 67.4 (2.6) cd | 50.4 (4.3) f | 0.53 (0.3) b | 69.2 b | 43.3 a | 7.8 (0.88) c |
| B73 | 30.26 (1.14) ab | 31.45 (2.19) a | 75.66 (5.22) b | 67.1 (1.61) cd | 1.21 (0.05) a | 86.59 a | 43.46 a | 14.34 (2.4) a |
| Mo17 | 24.6 (0.8) cde | 26.0 (0.9) cd | 63.0 (1.4) de | 66.1 (3.0) cde | 0.6 (0.2) b | 87.1 a | 53.7 a | 10.6 (2.1) abc |

**Fig. 3.**
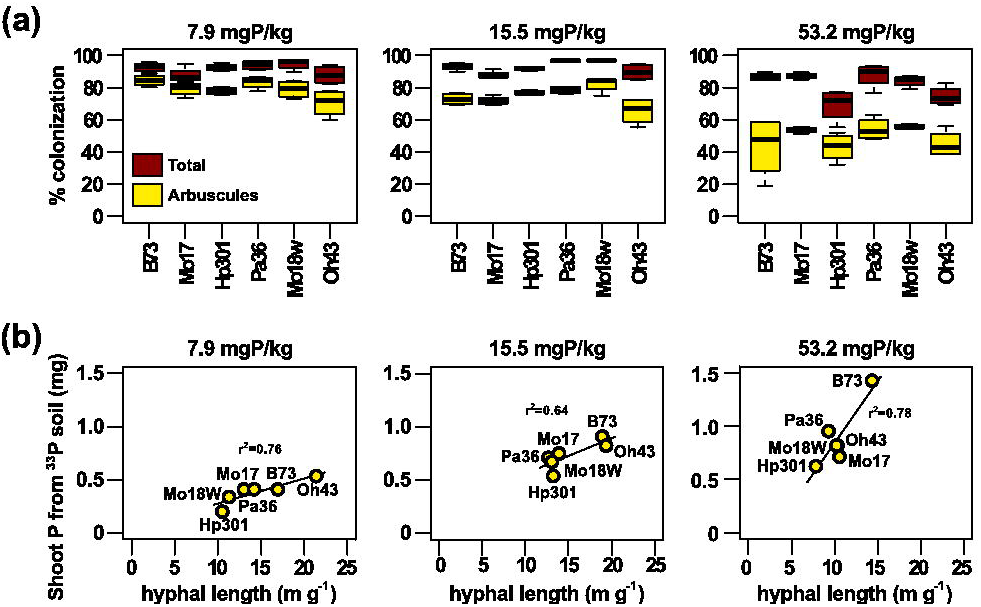
Phosphorus uptake from the hyphal compartment was correlated with the abundance of root-external hyphae in six maize lines inoculated with *Rhizophagus irregularis*. (a) Percentage of total root length containing mycorrhizal structures (brown) and arbuscules (yellow) in six maize lines, grown under three levels of phosphorus (P) availability (7.9, 15.5 and 53.2 mgP Kg^−1^) and inoculated with *R. irregularis.* Boxes show 1st quartile, median and 3rd quartile. Whiskers extend to the most extreme points within 1.5x box length. (b) Shoot P acquired from the hyphal compartment (Shoot P from ^33^P soil, mg) as a function of the length of root-external hyphal (m g^−1^ soil).

### Accumulation of *ZmPt* transcripts reflects functional differences among AM plants

To obtain further evidence that MR variation was linked to functional differences among M plants, accumulation of plant PHT1 phosphate transporter encoding transcripts (Fig. S3; Liu *et al*., 2016) was quantified in the roots of plants harvested from the *Rhizophagus irregularis* experiment. A preliminary analysis using the line B73 (Fig. S4) and previous reports (Fig. S6; Liu *et al*., 2016) were used to select the transcripts *ZmPt1*, *ZmPt3*, *ZmPt4*, *ZmPt5* and *ZmPt6* as being the most informative (based on tissue-specific expression and response to inoculation) for quantification in the six lines (Fig. 4). Across the six lines, the accumulation of *ZmPT* transcripts with respect to tissue type, inoculation with AM fungi and P availability was similar to that in B73 (Fig. S5). Within broad qualitative trends, however, quantitative differences were observed (Fig. 4).

**Fig. 4.**
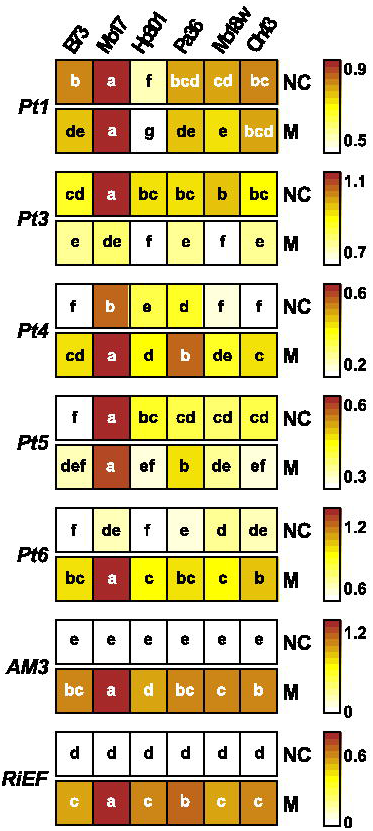
Accumulation of *ZmPt* transcripts responded in inoculation with *Rhizophagus irregularis.* Accumulation of six *ZmPt* (*Pt*) transcripts in root-samples of the lines B73, Mo17, Hp301, Pa36, Mo18W and Oh43, grown under low phosphorus (P) availability (7.9 mg kg^□1,^) with (M) or without (NC) inoculation with *R. irregularis*. Mean absolute transcript accumulation was determined from three biological replicates and scaled independently for each gene panel from white (minimum) to brown (maximum) accumulation. Letters indicate means groups based on least significant difference at α = 0.05, calculated independently for each gene. Accumulation of the maize marker transcript *ZmAm3* and the fungal transcript *RiEF* also shown.

To investigate patterns of variation in the accumulation of *ZmPt* transcripts, and to integrate transcript accumulation data with plant growth and physiological measurements, a principal component (PC) analysis was performed using the complete data set for M plants under low P (Fig. 5). In addition, pairwise correlations were calculated directly among all measurements (Fig. S5). The first two PCs captured 76% of the trait variation and well separated the six lines (Fig. 5). As expected from the previous analysis, plant dry weight was associated in the PC space with abundance of root-external hyphae, P uptake from the hyphae compartment (PHC) and shoot P content (Fig.5, lower right quadrant). Accumulation of *ZmPt1* and *ZmPt3* transcripts was associated with the same region of the PC space, and accumulation of *ZmPt3* transcripts was found to be positively correlated with root dry weight (RDW; p < 0.05), root P content (p < 0.1) and PHC (p < 0.1). Abundance of intra-radical fungal structures (%Col) was opposite to that of root-external hyphae in the PC space (Fig. 5, upper left compared with lower right quadrants), and a negative correlation was found between % Col and the length of root-external hyphae, suggesting a possible trade-off in the pattern of fungal growth. This correlation was, however, not significant at the 0.1 level indicating that this relationship did not hold for all lines. Accumulation of the marker transcripts *Am3* and *RiEF* was associated with accumulation of *ZmPt4*, *ZmPt5* and *ZmPt6*, and root P content (Fig. 5, upper right quadrant). Interestingly, although accumulation of *ZmPt6* strongly differentiated NC and M plants (Fig. S5), among these six lines, when inoculated, no correlation was observed between *ZmPt6* accumulation and arbuscule abundance. Placing the lines on the PC space, Oh43 was separated from other lines on the basis of PC2 (Fig. 5, lower right quadrant), associated with shoot P, length of root-external hyphae, PHC and to a lesser extent accumulation of *ZmPt4* and *ZmPt5.* The lines Mo18W and Hp301 were associated with low biomass and low levels of *ZmPt* transcript accumulation, but high levels of intra-radical colonization (Fig. 5, upper left quadrant). Mo17 was distinct in accumulating high levels of AM associated transcripts, although with no associated increase in colonization, development of root-external hyphae or MR. Taken as a whole, physiological and molecular data support the interpretation that superior MR in Oh43 results from the nature of the fungal-plant interaction, and is not an artifact resulting from a high level of dependence.

**Fig. 5.**
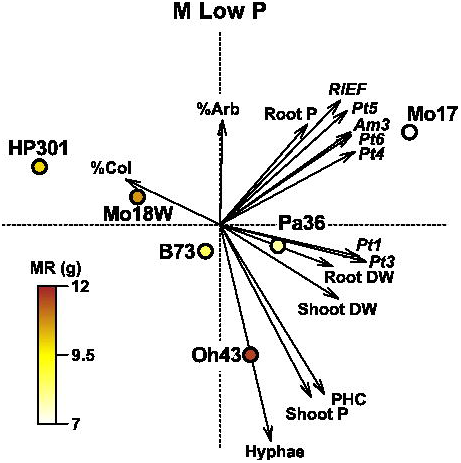
Molecular and physiological patterns were correlated in maize plants inoculated with *Rhizophagus irregularis.* Principle component analysis (PCA) of plant-growth, physiological and molecular observations of six maize lines grown under low phosphorus (P) availability (7.9 mg kg^□1^) and inoculated with *R. irregularis*. Biplot showing scores in the first two principal components (PC1: x-axis, PC2: y-axis) for traits (black arrows: shoot dry weight (shoot DW), shoot P, root dry weight (root DW), root P, total colonization (%Col), arbuscule abundance (%Arb), length of root-external hyphae (Hyphae), P uptake from the hyphal compartment (PHC), accumulation of *Pt1*, *Pt3, Pt4*, *Pt5*, *Pt6*, *Am3* and *RiEF* transcripts) and lines (B73, Mo17, HP301, Pa36, Mo18W, Oh43). Points indicating the different lines are coloured by mycorrhizal response (MR, g) calculated as the difference in shoot dry weight in colonized and non-colonized plants.

## DISCUSSION

Data presented in this study reveal genetic variation in the capacity of maize varieties to profit from AM symbiosis, beyond differences in plant dependence (Janos 2007; Sawers *et al.*, 2008, 2010). Evaluation of the relative growth of thirty highly diverse lines (McMullen *et al.*, 2009) with and without inoculation with *Funneliformis mosseae* distinguished those that were highly responsive on the basis of poor performance in the absence of symbiosis (*i.e.* highly dependent) from those that benefited more from the symbiosis *per se*. The line Oh43 was selected as an example of the latter. Support for this initial interpretation was obtained by detailed physiological and molecular characterization linking superior responsiveness of Oh43 to *Rhizophagus irregularis* with enhanced PAE in mycorrhizal plants and a greater abundance of root-external hyphae.

Phosphorus limitation of plant growth results primarily from the low mobility of P in the soil. In this study, it was observed that increased plant growth in AM colonized plants was accompanied by greater shoot P content. Once P has reached the root-surface or been delivered to the peri-arbuscular space, subsequent uptake by plant PHT1 transporters is not predicted to limit PAE (Bucher, 2007). As is consistent, it was neither the extent of intra-radical colonization nor the accumulation of *ZmPt6* transcripts (predicted to encode the major peri-arbuscular membrane associated PT transporter) that showed the greatest correlation with P uptake, but the abundance of root-external hyphae. This generally supports previous studies (*e.g.* Schweiger & Jakobsen 1999, Jakobsen *et al.*, 2001, Yao *et al.*, 2001, Schnepf *et al.*, 2008), including an evaluation of diverse fungal isolates used to inoculate a common plant host, which found a similar correlation between fungal P uptake and hyphal length (Munkvold *et al.*, 2004). Although a further report characterizing variation in mycorrhiza response among four Chinese maize varieties did not reveal a clear relationship between P uptake and the length of root-external hyphae (Chu *et al.*, 2013), this may reflect the specific genotypes evaluated, or the fact that the contribution of mycorrhizal P uptake was not directly quantified. A potential trade-off was observed between the abundance of intra-radical and root-external fungal structures, the balance of which is apparently influenced by plant genetic factors. Given the importance of hyphal abundance to PAE, the data suggest that quantification of intra-radical structures alone is not predictive of P uptake via the mycorrhizal pathway or growth response.

Prior physiological characterization has demonstrated that mycorrhizal P uptake is not a simple addition to direct P uptake, but may represent a functional alternative: in the extreme case, a colonized plant may obtain nearly all of its P requirement via the AM pathway, whether as a result of down regulation of the direct pathway or owing to a greater efficiency of fungal P foraging compared with that of plant roots (Smith *et al.*, 2003; Schnepf *et al.*, 2008). Furthermore, the mycorrhizal pathway may remain important at higher P availability, even when MR itself is small, or even negative. Following inoculation with *R. irregularis*, the absolute quantity of P obtained via the AM pathway was greater at high P than at low P, even though the abundance of intra-radical and root external fungal structures was lower, presumably as the result of increased Pi in solution as the capacity of the soil to adsorb P was saturated. Collectively, these data illustrate the complexity of determining symbiotic outcome, and the range of plasticity among just six lines, with respect to just a single environmental variable.

Transcriptome profiling has identified rice marker genes whose transcript accumulation correlates well with the establishment and development of AM symbiosis, differentiating colonized from non-colonized plants (*e.g.* Guimil *et al*., 2005; Gutjahr *et al*., 2015). In this study, transcripts encoding PHT1 phosphate transporters were quantified primarily not to distinguish NC and M plants in general, but to investigate differences among the different lines when colonized. Overall, AM colonization was positively correlated with the accumulation of transcripts encoded by *Pt6*, and to a lesser extent those encoded by *Pt4, Pt5* and *Pt2*, consistent with previous reports (Nagy *et al.*, 2006; Willmann *et al.*, 2013; Liu *et al*., 2016). Variation in the accumulation of the *ZmPt6* transcript, however, was largely independent of differences in growth response to *R. irregularis*, although arbuscule abundance itself was also a poor predictor of MR among the six lines evaluated in this study. These observations are consistent with the interpretation that P transfer at the peri-arbuscular interface is non-limiting, with respect to either arbuscule abundance or the concentration of PT6 proteins in the peri-arbuscular membrane. PT6 protein, however, has been reported also to regulate developmental responses to P limitation (Volpe *et al*., 2016), and variation in *ZmPt6* accumulation may have additional significance beyond P transfer. A number of additional maize *ZmPt* transcripts responded to AM inoculation, although they were less abundant than those encoded by *ZmPt6.* Significantly, at low P, a mild positive correlation was observed between accumulation of *ZmPt4* and *ZmPt5* transcripts and shoot biomass among colonized plants, indicating a role in the regulation of the symbiosis. Accumulation of *Pt1* and *Pt3* transcripts, although generally lower in M than NC plants, was positively correlated with root and shoot dry weight among M lines. Accumulation of *ZmPt1* was positively correlated also with P uptake from the hyphal compartment. Interestingly, the correlation was stronger with dry weight than P content, indicating this may be more than a secondary effect of differences in P accumulation. These observations, along with previous characterization of *pt11* and *pt13* mutants in rice (Yang *et al*., 2012), suggest a role for PHT1 proteins not only in P uptake but in the fine tuning of cost-benefit in AM symbioses.

Previous characterization of variation in MR has placed an emphasis on the development of intra-radical fungal structures, and marker transcripts have been identified allowing molecular-based quantification of intra-radical colonization. In this study, it was observed that variation in the abundance of root-external hyphae was better correlated than levels of intra-radical colonization with plant growth response to inoculation with *R. irregularis*. Although accumulation of the well characterized *ZmPt6* transcript was not predictive of the abundance of extra-radical hyphae, correlations were observed between transcripts encoded by other *ZmPt* genes, abundance of extra-radical hyphae and mycorrhiza response. The identification of such variation, coupled with the availability of populations for quantitative trait loci mapping (McMullen *et al.*, 2009), opens up the possibility to characterize the genetic basis of host effects on the development of extra-radical hyphae, and to develop molecular breeding strategies to target this important, but hard to evaluate, component of the outcome of mycorrhizal symbiosis.

## ACKNOWLEDGEMENTS

This work was supported by the Swiss National Science Foundation ‘professeur boursier’ grants PP00A-110874, PP00P3-130704, by the Gatsby Charitable Foundation grant RG60824, by The Danish Council for Independent Research, Technology and Production Sciences grant 0602-01412B and by the Mexican National Council of Science and Technology (CONACYT) grant CB2012-151947. We thank Mesfin Nigussie Gebreselassie and Matthias Mueller for assistance with plant growth and evaluation.

## AUTHOR CONTRIBUTION

Diversity screen: RS, BW, UP. Ionomic analysis: IB. Physiological characterization of phosphate uptake in selected lines: SS, MG, IJ. Identification of *Pht1* genes and phylogenetic analysis: EGM, RCM, RS. Quantification of transcript accumulation: CQ, SS. Statistical analysis: JG, RS. Experimental design: UP, IJ, RS. All authors contributed to data interpretation and writing of the manuscript.

## SUPPORTING INFORMATION

The following Supporting Information is available for this article:

**Fig. S1** Inoculation with *Funneliformis mosseae* promotes growth in a panel of 30 diverse maize lines under low phosphorus availability

**Fig. S2** PHT1 protein tree

**Fig. S3** Maize *PHT1* transcripts accumulate throughout the plant life-cycle

**Fig. S4** Accumulation of transcripts encoding PHT1 proteins responds to phosphorus availability and inoculation with *Rhizophagus irregularis*.

**Fig. S5** Accumulation of transcripts encoding PHT1 proteins is correlated with phosphorus accumulation and inoculation with *Rhizophagus irregularis.*
across diverse maize lines.

**Table S1** Phosphorus concentration in the leaves of 30 maize lines grown with or without inoculation with *Funneliformis mosseae*.

**Table S2** Primers used for real time PCR

